# Multivalent Tri-Functional T-cell Engagers by Chemically Induced Protein Self-Assembly

**DOI:** 10.1101/2025.07.29.667528

**Authors:** Declan Dahlberg, Brandi McKnight, Abhishek Kulkarni, Lakmal Rozumalski, Freddys Rodriguez, Yiao Wang, Carston R. Wagner

**Author notes:** Address correspondence to, University of Minnesota, Department of Medicinal Chemistry, 2231 6th Street S.E., Cancer & Cardiovascular Research Building, Minneapolis, Minnesota 55455, USA. Contributed equally (Brandi McKnight, Abhishek Kulkarni).

## Abstract

T-cell engagers (TCEs) show promise in cancer immunotherapy but face challenges in solid tumors due to heterogeneity, antigen escape, and limited T-cell infiltration. To address this, we developed a modular platform using chemically self-assembled nanorings (CSANs). We engineered a bifunctional fusion protein, E1-DHFR^2^-αCD3, with an EGFR-binding fibronectin (E1) and an anti-CD3 scFv on a DHFR^2^ scaffold. With bis-methotrexate, the monomers formed multivalent cis-CSANs.

Both monomers and CSANs bound EGFR+ tumor cells and T-cells, were internalized, and induced dose-dependent, EGFR- and T-cell-dependent cytotoxicity in co-culture assays. The system was reproducible across T-cell donors. To expand targeting, E1-DHFR2-αCD3 co-assembled with other DHFR2 monomers targeting GFP or EpCAM, forming trispecific CSANs capable of binding multiple antigens and mediating cytotoxicity.

This platform enables the discovery of potent multispecific TCEs to address antigen escape and solid tumor heterogeneity. Future work will focus on optimizing antigen combinations to enhance efficacy across breast cancer subtypes.

## Introduction

The utilization of methods to target T-cells to cancer cells, such as CAR (chimeric antigen receptor) T-cells, has rapidly become an arm of the cancer immunotherapy armamentarium.^1–3^ Alternative approaches, such as bispecific antibodies and immune cell engagers, that do not require genetic modification of a patient’s T-cells, have emerged as an attractive off-the-shelf alternative.^4^ This class of therapeutics, broadly termed immune cell engagers, utilize various scaffolds to non-genetically modify T-cells using a T-cell binder (e.g., αCD3) and redirect them to malignant tissue with an effector arm of choice (e.g., αCD19, αEGFR, αEpCAM, αPSMA, etc.).^5^ While these bispecific constructs were met with initial enthusiasm, targeting only a single receptor on the surface of immune and cancer cells presents significant challenges, including tumor-intrinsic and tumor-extrinsic resistance mechanisms.^6^ As such, engineering multispecific constructs to overcome these challenges is an attractive consideration.^7,8^

Our laboratory has developed a novel immune cell engager from chemically self-assembled nanorings (CSANs). CSANs are formed utilizing a fusion of two *E. coli* dihydrofolate reductase proteins (DHFR^2^) which can be incubated with the molecular glue, bis-methotrexate (bis-MTX), to form multivalent rings. By modifying either terminus of DHFR^2^ with a T-cell or tumor targeting element, bispecific and multivalent rings can be formed. The multivalency of CSANs creates an enhanced avidity effect and a high degree of selectivity to the cells of interest.^9^ Previously, bispecific T-cell engaging CSANs have been prepared by mixing a DHFR^2^-αCD3 fusion protein with another DHFR^2^ fusion protein incorporating a tumor targeting ligand.^10–12^ These CSANs are referred to as *trans-*CSANs because they are made of two different monospecific DHFR^2^ proteins **(Figure 1A)**. We have shown the versatility of this approach by utilizing the DHFR^2^-αCD3 monomer to prepare bispecific CSANs with tumor targeting single-chain variable fragments (scFvs) and fibronectins.^10–13^ We hypothesized that bispecific *cis-*CSANs **(Figure 1B)** could be constructed by modifying both termini of the DHFR^2^ scaffold to produce bispecific monomers, which would provide a path for the preparation of trispecific, multivalent CSANs when combined with a DHFR^2^ monomer **(Figure 1C)**. To address our hypothesis, we designed and prepared DHFR^2^ proteins incorporating the epidermal growth factor receptor (EGFR) binding fibronectin (E1),^14^ and αCD3 scFv [herein referred to as E1-DHFR^2^-αCD3]. We demonstrate that the bispecific *cis-*CSANs construct functions as a potent multivalent T-cell engager, resulting in highly stable interactions between T-cells and the surface exposed target antigen, EGFR. In addition, we demonstrate with the bispecific *cis-*CSANs that trispecific CSAN T-cell engagers can easily be prepared **(Figure 1C-D)**.

**Figure 1.**
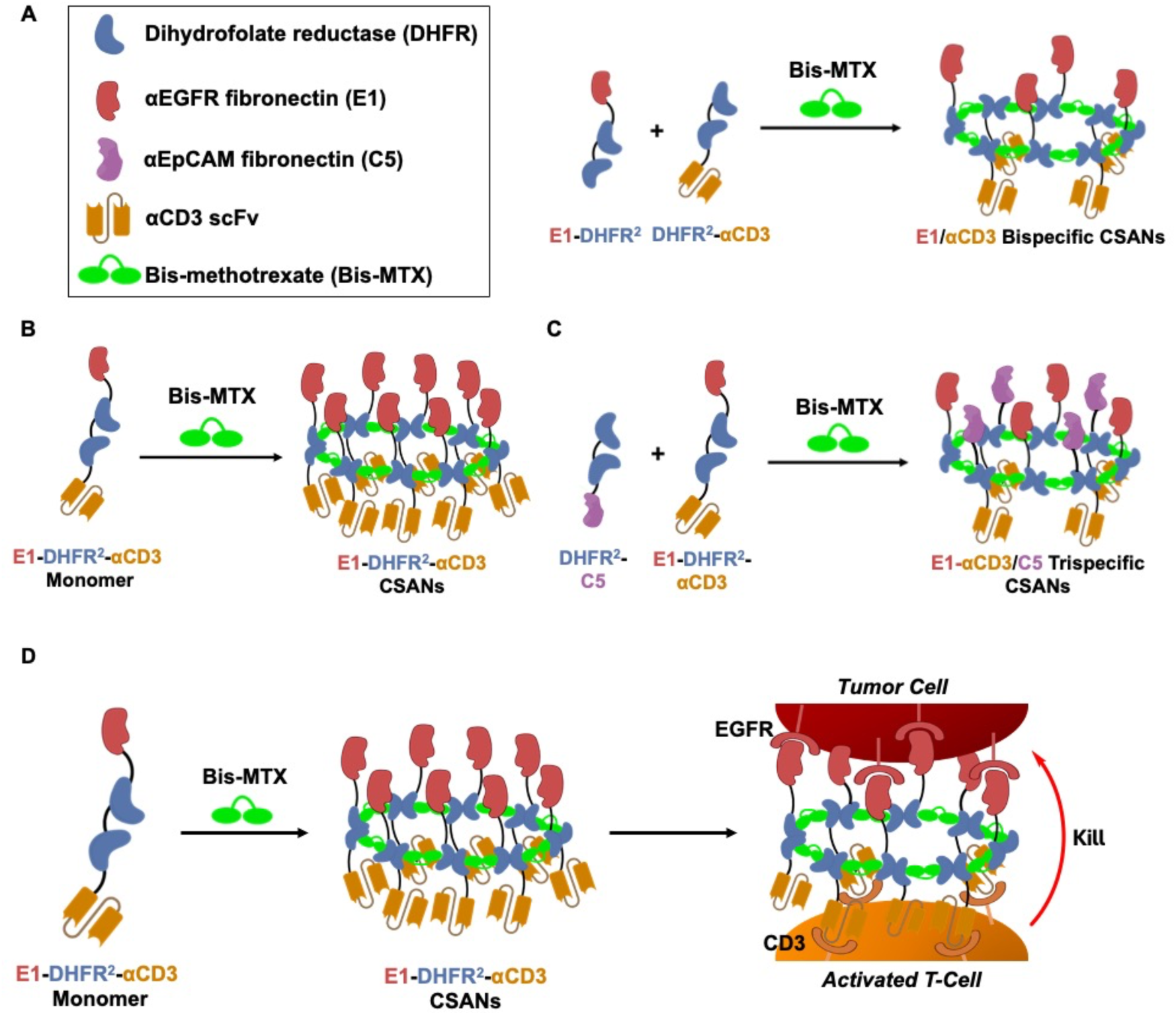
Structures of EGFR and CD3 targeted chemically self-assembled nanorings (CSANs). **A)** αEGFR/αCD3 *trans*-CSANs prepared from DHFR^2^-αCD3 and E1-DHFR^2^ fusion proteins. **B)** αEGFR-αCD3 *cis*-CSANs prepared from E1-DHFR^2^-αCD3 bispecific fusion protein. **C)** αEGFR-αCD3/αEpCAM trispecific CSANs prepared from E1-DHFR^2^-αCD3 and DHFR^2^-αEpCAM (C5) fusion protein. **D)** Proposed mechanism of *cis-*CSAN mediated, T-cell directed killing of EGFR+ tumor cells.

## Results

### Preparation and Characterization of E1-DHFR^2^-αCD3 Fusion Protein

The E1-DHFR^2^-αCD3 fusion protein is composed of two targeting elements fused to our DHFR^2^ scaffold **(Figure 1A)**. In the sequence design, we attached an αEGFR fibronectin named E1, previously utilized to prepare E1-DHFR^2^, to the N-terminus and an αCD3 scFv, previously characterized in our lab, to the C-terminus **(Figure S1A)**. Additionally, the sequences included a FLAG tag at the N-terminus and a hexa-histidine tag (His6) at the C-terminus. The gene for E1-DHFR^2^-αCD3 was cloned into the pET28a vector, transformed into T7® express *E coli*., and then expressed as an insoluble protein. E1-DHFR^2^-αCD3 protein was then purified from inclusion bodies, and its purity was demonstrated by SDS-PAGE. **(Figure S1B)**. To ensure the correct protein was obtained, E1-DHFR^2^-αCD3 was analyzed using liquid chromatography-mass spectrometry (LC-MS) and compared to the theoretical molecular weight (MW) of 78,550 Da. Analysis of the LC-MS data showed a MW of 78,570 Da **(Figure 2A)**, which closely matches our theoretical MW ensuring that we were able to properly express and purify E1-DHFR^2^-αCD3.

**Figure 2.**
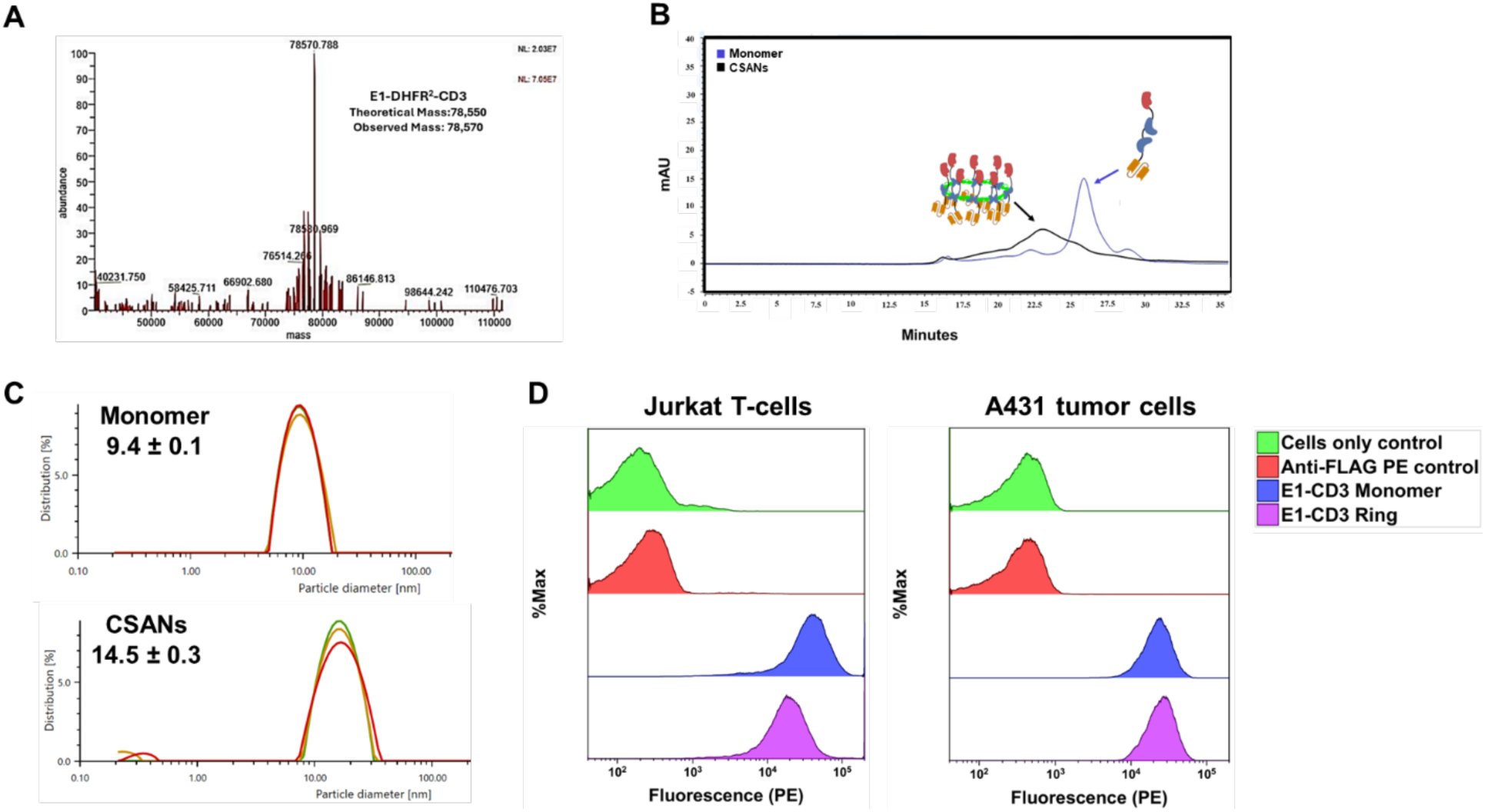
Characterization of E1-DHFR^2^-αCD3 protein. **A)** LC-MS demonstrating E1-DHFR^2^-αCD3 protein matches theoretical molecular weight. **B)** SEC analysis of E1-DHFR^2^-αCD3 monomer and CSANs after 1 hour incubation with bis-MTX. **C)** Dynamic light scattering showing hydrodynamic diameter of E1-DHFR^2^-αCD3 monomer and CSANs. **D)** Flow cytometry showing bifunctionality of E1-DHFR^2^-αCD3, and its ability to bind Jurkat T-cells via CD3 and A431-R cells via EGFR as a monomer and CSAN.

After expressing and purifying E1-DHFR^2^-αCD3, analytical size exclusion chromatography (SEC) was used to assess protein purity and CSAN formation after incubation with bis-MTX for 1 hour at room temperature **(Figure 2B)**. E1-DHFR^2^-αCD3 monomer was then compared to E1-DHFR^2^ and DHFR^2^-αCD3 monomeric proteins on SEC. E1-DHFR^2^-αCD3 (78.5 kDa) exhibited a larger size, eluting earlier compared to both E1-DHFR^2^ (48.7 kDa) and DHFR^2^-αCD3 (64 kDa) **(Figure S1C)**. Previously, we have demonstrated that typical *trans*-CSANs are composed of, on average, eight monomers.^15^ Given the increased size of the E1-DHFR^2^-αCD3 monomer relative to the E1-DHFR^2^ and DHFR^2^-αCD3 monomers, we expected the corresponding CSANs to be of greater molecular weight and hydrodynamic diameter. Surprisingly, SEC analysis of *cis*-CSANs formed by E1-DHFR^2^-αCD3 were found to elute later than E1/αCD3 *trans*-CSANs formed from E1-DHFR^2^ and DHFR^2^-αCD3 monomers, indicating a potentially smaller hydrodynamic diameter **(Figure S1D)**. Consistent with the SEC analysis, dynamic light scattering (DLS) studies revealed that while, as expected, the E1-DHFR^2^-αCD3 monomer hydrodynamic diameter (Dh) was larger than either E1-DHFR^2^ and DHFR^2^-αCD3 monomer, (Dh = 9.36 ± 0.1 nm vs 6-7 nm), the E1-DHFR^2^-αCD3 *cis*-CSANs exhibited a smaller Dh value than the corresponding E1/αCD3 *trans*-CSANs, 14.48 ± 0.27 nM and 18-20 nm, respectively (**Figure 2C/S1E**). Thus, the size obtained for the E1-DHFR^2^-αCD3 *cis*-CSANs was smaller than what has been observed for our traditional E1/αCD3 *trans*-CSANs, consistent with the results seen in our SEC data. To verify that the αEGFR (E1) and αCD3 domains of E1-DHFR^2^-αCD3 were functional, flow cytometry experiments were performed with Jurkat T-cells and the red fluorescent protein mKate2 expressing epidermoid carcinoma cell line A431-R (EGFR+, 5.8 ×10^6^ per cell). Using the N-terminal FLAG tag for detection, binding was observed for both monomer and CSAN forms of αEGFR (E1)-DHFR^2^-αCD3 to Jurkat T-cells and A431-R cells **(Figure 2D)**. Similarly, the αEGFR (E1)-DHFR^2^-αCD3 monomer and corresponding CSANs were also found to bind to normal human T-cells and the green fluorescent protein (GFP) expressing triple negative breast cancer (TNBC) cell line, MDA-MB-231-G, which express approximately 100-fold (6.8 × 10^4^ per cell) less EGFR than A431-R cells **(Figure S2A-B)**.

When EGFR is activated through ligand binding, the receptor is internalized through endocytosis and then trafficked to the early endosomal compartment of the cell. Once internalized, the receptor can either be degraded or recycled to the cell surface.^16^ Previous work in our lab revealed E1/αCD3 *trans*-bispecific CSANs exhibit EGFR-dependent endocytosis, which is temperature dependent, suggesting energy dependent endocytosis.^11^ To assess the ability of the E1-DHFR^2^-αCD3 monomer and the corresponding CSANs to undergo internalization, E1-DHFR^2^-αCD3 was labeled nonspecifically with an Alexa Fluor 647 NHS ester and then assembled into CSANs by incubation with bis-MTX. DAPI-stained MDA-MB-231-G cells were treated for 1 hour at 37 ⁰C, with either the Alexa Fluor 647-labelled E1-DHFR^2^-αCD3 monomer or corresponding CSANs, followed by imaging. Internalization of both the monomeric and CSAN forms of E1-DHFR^2^-αCD3 was observed, as indicated by internal punctates and localization of the labeled protein near the cell nuclei. In contrast, cellular internalization of the DHFR^2^-αCD3 CSANs, which lack the αEGFR-fibronectin (E1), was not observed. In addition, internalization of the E1-DHFR^2^-αCD3 monomer and CSANs was not observed at 4 °C, consistent with previous results demonstrating that the αEGFR-fibronectin (E1) induces temperature dependent, EGFR-mediated endocytosis^11^ **(Figure 3A-B)**.

**Figure 3.**
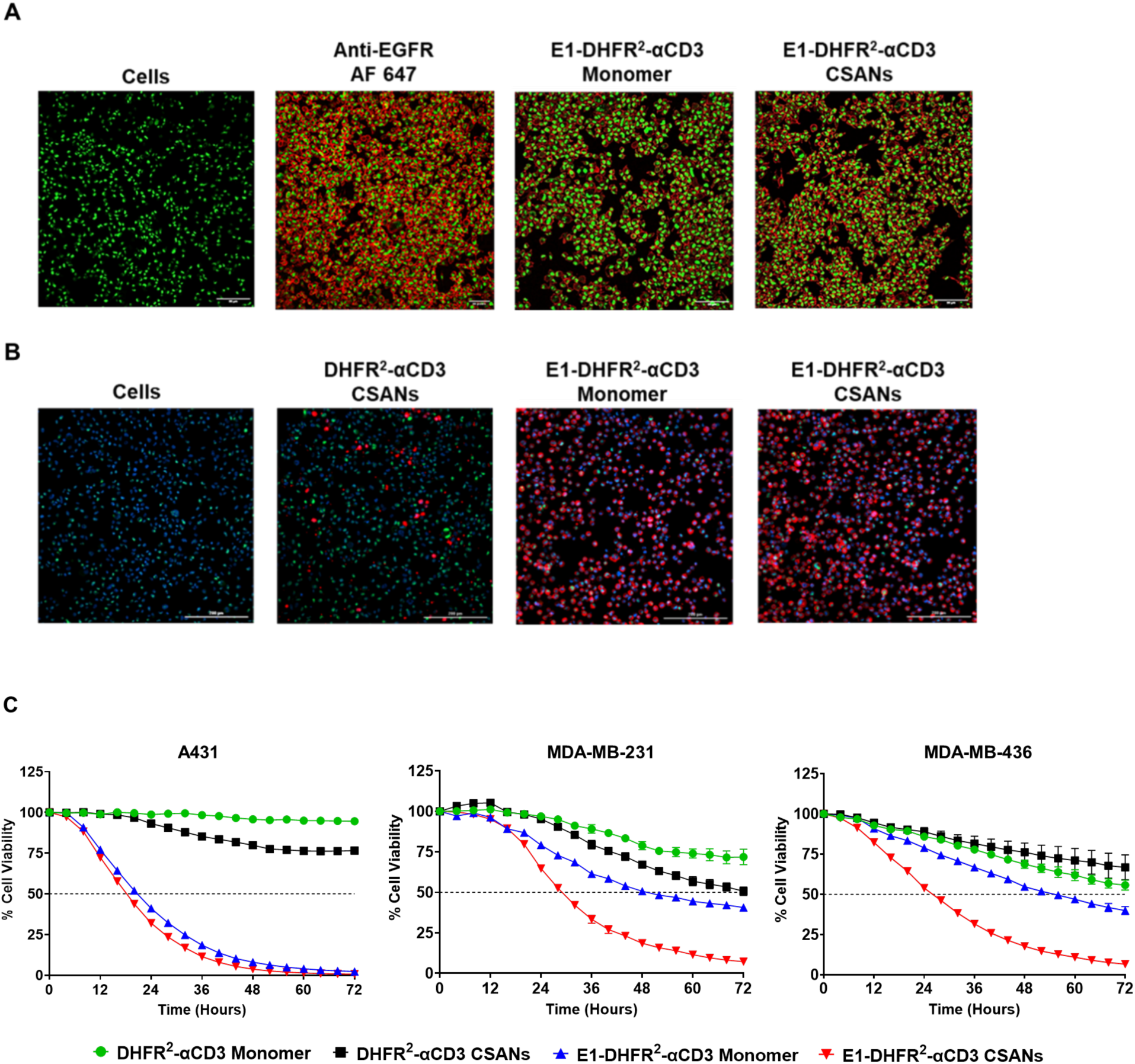
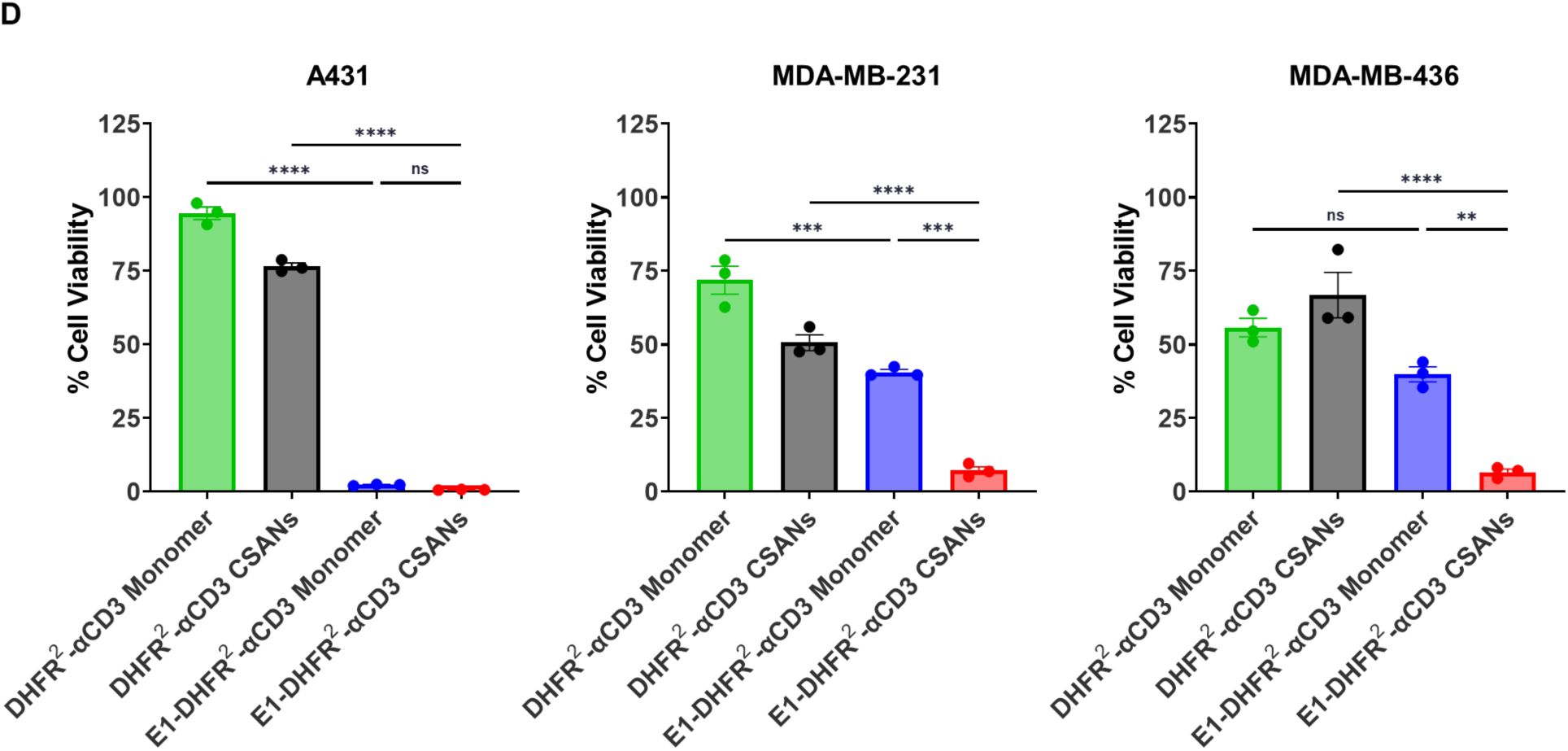
Internalization of E1-DHFR^2^-αCD3 by MDA-MB-231-G cells and cytotoxicity verification across multiple EGFR+ cell lines, including TNBC. A monolayer of nuclear-stained (blue) MDA-MB-231-G cells (green) was incubated with either PBS, Alexa Fluor 647-labelled DHFR^2^-αCD3 CSANs, E1-DHFR^2^-αCD3 monomer, or E1-DHFR^2^-αCD3 CSANs and imaged after 1 hour of incubation at **A)** 4 °C and **B)** 37 ⁰C. Post-incubation, it was observed that both the E1-DHFR^2^-αCD3 monomer and CSANs were internalized only at 37 °C, as evidenced by the presence of the proteins (red) near the cell nuclei (blue). Samples incubated at 4 °C remained bound to the surface of the cells and did not demonstrate internalization. **C)** A431-R, MDA-MB-231-G, or MDA-MB-436-R cells were seeded and cultured as a monolayer in a 96-well plate. T-cells isolated from donor-derived PBMCs were incubated with either monomeric protein or CSANs (25 nM) for an hour to form prosthetic antigen receptor (PAR) T-cells. These PAR T-cells were added to the target cells at a ratio of 10:1 (Effector:Target) ∼16 hours after seeding the cancer cells. Tumor cell viability was continuously monitored over 72 hours using the BioTek Biospa Live Cell Imaging Analysis System. Data are represented as mean ± SEM of n=3 replicates. **D)** Endpoint quantification of cytotoxicity. Each column represents the final % target cell viability at the end of the study. Data are presented as mean ± SEM of n=3 replicates with statistical significance denoted as *P<0.05, **P<0.01, ***P<0.001, and ****P<0.0001 by one-way ordinary ANOVA followed by Tukey’s multiple comparisons test. Data obtained from T-cells isolated from a single PBMC donor but is representative of two different donors.

### E1- DHFR^2^-αCD3 *cis-*CSANs Facilitate Cytotoxicity Against Multiple EGFR+ Cell Lines Including Triple-Negative Breast Cancer Cells

The ability of the E1-DHFR^2^-αCD3 monomer and corresponding *cis-*CSANs to induce T-cell mediated tumor cell cytotoxicity was interrogated using multiple cell lines expressing various amounts of EGFR [transduced with either mKate-2 (-R) or GFP (-G)]. A431-R cells were considered “EGFR+ high-expression” while the TNBC cell lines MDA-MB-231-G and MDA-MB-436-R were considered “EGFR+ low-expression” as quantified using Bangs Laboratory Calibration Beads **(Table S1)**. T-cells isolated from donor-derived peripheral blood mononuclear cells (PBMCs), were incubated with 25 nM of the E1-DHFR^2^-αCD3 monomer and *cis*-CSANs, and the cytotoxicity was determined over 72 hours **(Figure 3C)**. After quantifying endpoint cytotoxicity, we demonstrate that the E1-DHFR^2^-αCD3 construct induces potent cytotoxicity against all three cell lines both in monomeric and CSANs form **(Figure 3D)**. Interestingly, there was no statistical significance between monomer and CSANs for A431-R cells, suggesting that with high surface expression of EGFR, the monomeric E1-DHFR^2^-αCD3 construct can induce equipotent target cell lysis to the multivalent CSANs. Conversely, we demonstrate that the CSANs perform better when EGFR expression is low (MDA-MB-231-G and MDA-MB-436-R) than monomeric E1-DHFR^2^-αCD3. These assays were repeated with multiple effector-to-target (E:T) ratios, which may better represent the immune cell population in the tumor microenvironment for both A431-R cells **(Figure 4A)** and MDA-MB-231-G cells **(Figure S3A)**.

**Figure 4.**
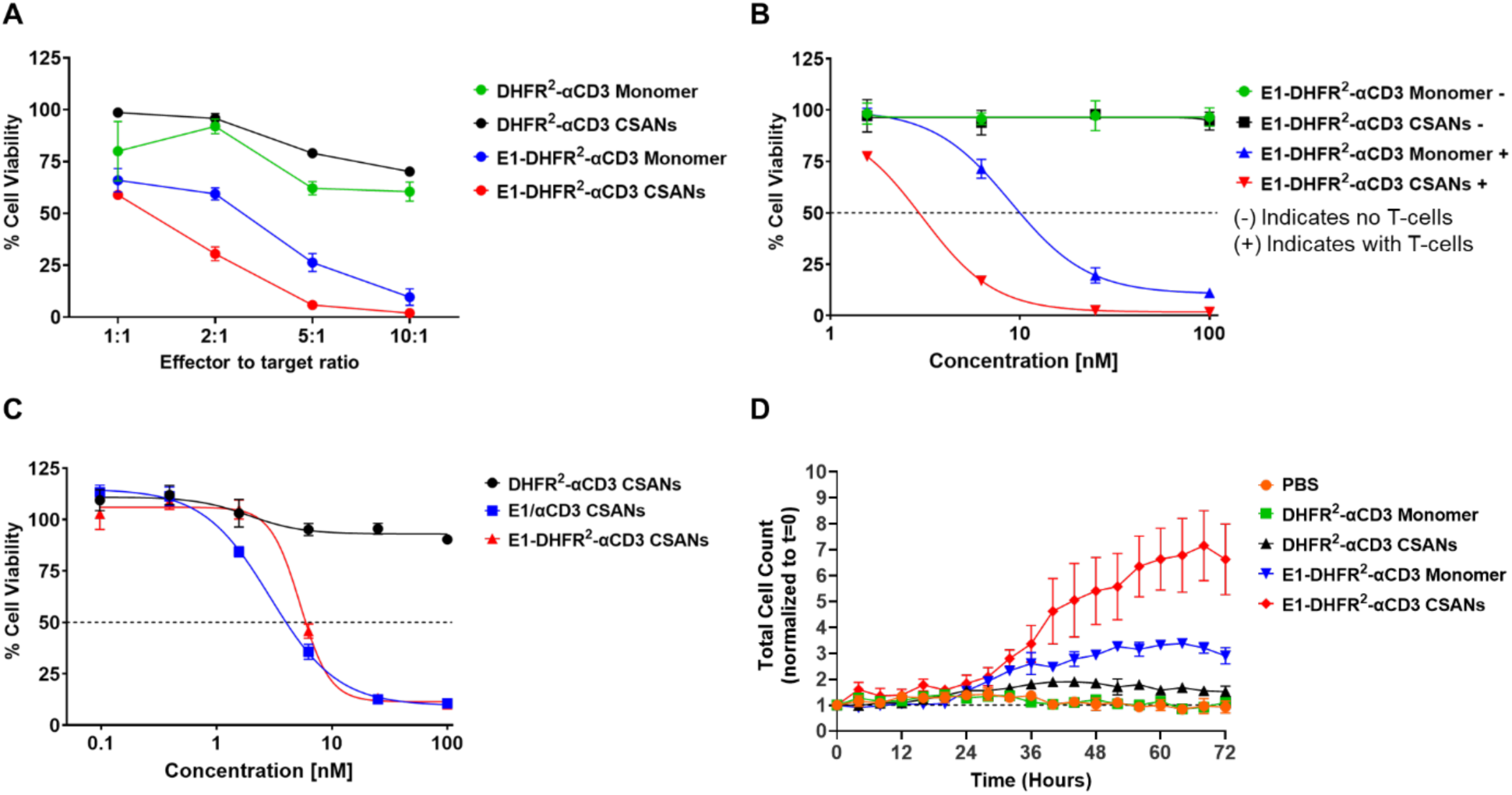
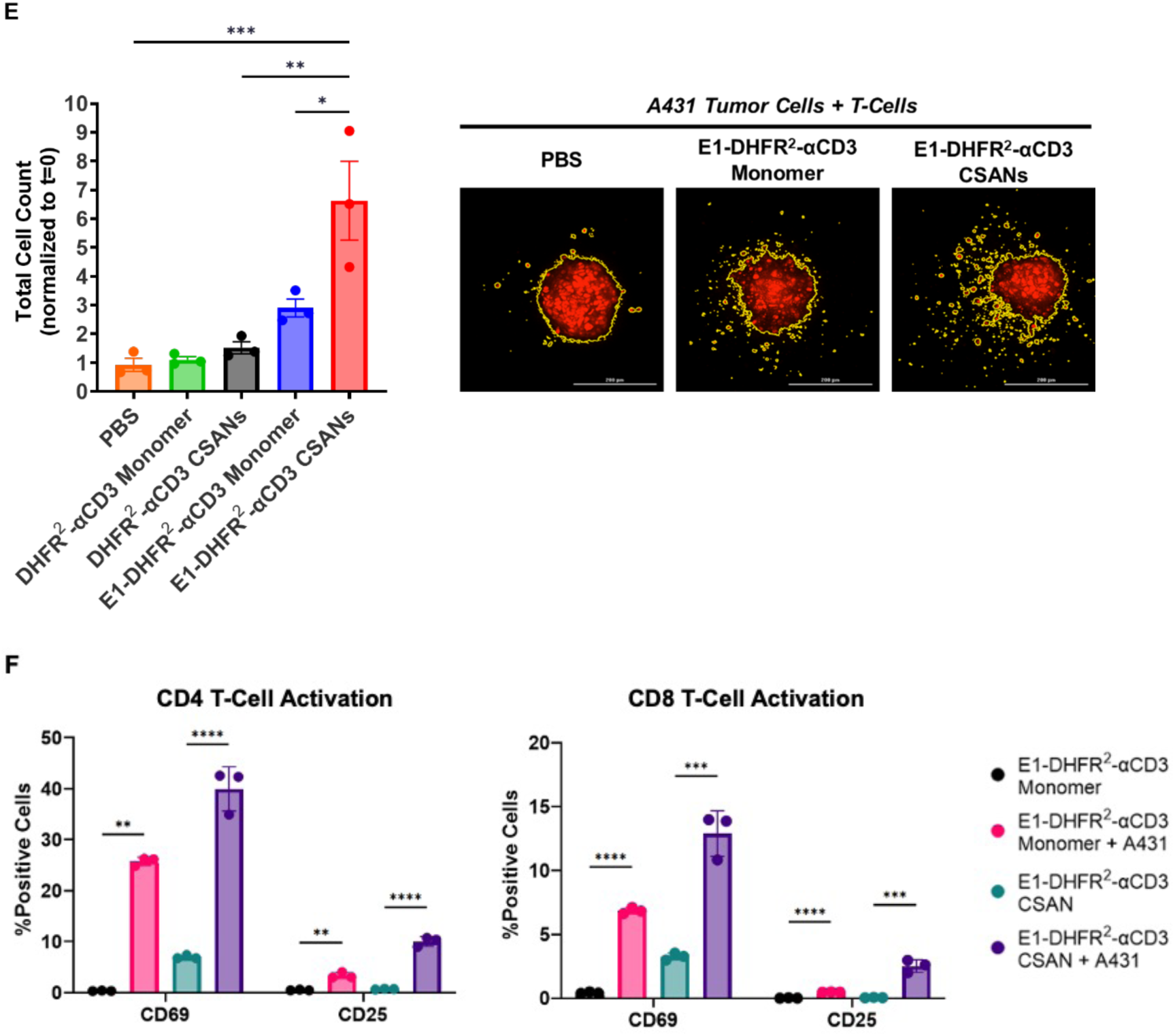
E1-DHFR^2^-αCD3 exhibits comparable cytotoxicity to *trans-*CSANs, maintains cytotoxicity in 3D culture, and selectively activates T-cells. **A)** Target cells and PAR T-cells were co-cultured as outlined previously. A431-R cell viability was tracked over 72 hours using different E:T ratios (1:1, 2:1, 5:1, 10:1) and a constant concentration of monomer and CSANs at 25 nM. Data points represent % target cell viability at the conclusion of the 72-hour study. Data are presented as mean ± SEM of n=3 replicates. **B)** Target cells and PAR T-cells were co-cultured as outlined previously. A431-R cell viability was tracked over 72 hours using different monomer and CSAN concentrations, either with (E:T of 10:1) or without T-cells. Data points represent % target cell viability after the 72-hour study. Data are presented as mean ± SEM of n=3 replicates. **C)** E1-DHFR^2^-αCD3 *cis*-CSANs, and E1/αCD3 *trans-*CSANs were cultured with A431-R target cells and PAR T-cells as outlined previously using multiple concentrations. Endpoint data was plotted and analyzed with *Prism* dose-response IC_50_ curve fitting to yield IC_50_ ranges with a 95% confidence interval. **D)** 5000 A431-R cells were centrifuged in a 96-well ultra-low attachment (ULA) round-bottom plate, and spheroids were allowed to form overnight. PAR T-cells were then added to plates and spheroids were imaged every 4 hours. After 72 hours, Z-projections of spheroids were compiled and time-lapse was used to show spheroid disruption. Cell counting function (yellow mask in the image) was then used to track and plot the increase in cell count (disruption) over time. **E)** Endpoint comparison at 72 hours analyzed where data are presented as mean ± SEM of n=3 replicates with statistical significance denoted as *P<0.05, **P<0.01, ***P<0.001, and ****P<0.0001 with one-way ordinary ANOVA followed by Tukey’s multiple comparisons test. **F)** T-cell activation was monitored by flow cytometry after a 24-hour cytotoxicity assay. 25 nM of E1-DHFR^2^-αCD3 monomer or CSANs were incubated with either T-cells alone or T-cells and A431-R target cells at an E:T ratio of 10:1. Expression of CD69 and CD25 was measured by flow cytometry on both CD4+ and CD8+ T-cells from all experimental groups. Samples were run on the Cytoflex-S flow cytometer and analyzed with Kaluza software. Data are presented as mean ± SEM of n=3 replicates with statistical significance denoted as *P<0.05, **P<0.01, ***P<0.001, and ****P<0.0001 by two-way ANOVA followed by Dunnett’s test for multiple comparisons. Data obtained from T-cells from a single PBMC donor.

### Quantitative T-Cell Dependent Cytotoxicity Comparison of E1-DHFR^2^-αCD3 *Cis-*CSANs to E1/αCD3 *Trans*-CSANs

To quantify the potency of E1-DHFR^2^-αCD3 *cis*-CSANs we first demonstrated that both E1-DHFR^2^-αCD3 monomer and CSANs exhibit potent cytotoxicity across a range of low nanomolar concentrations with a fixed effector to target ratio (10:1) and that the proteins do not exhibit background cytotoxicity without T-cells present **(Figure 4B)**. Next, to demonstrate that the incorporation of the E1 and αCD3 targeting ligands on a single scaffold do not affect the cytotoxicity compared to previous work with *trans-*CSANs, we performed an IC_50_ analysis using E1-DHFR^2^-αCD3 *cis-*CSANs, E1/αCD3 *trans-*CSANs, and DHFR^2^-αCD3 CSANs as a control. The calculated IC_50_ for the *cis* and *trans* constructs were 5.74 and 2.89 nM respectively **(Figure 4C)**. DHFR^2^-αCD3 CSANs used as a control yielded IC_50_ >100 nM.

### E1-DHFR^2^-αCD3 CSANs are Cytotoxic in an *In-Vitro* 3D Spheroid Model and Selectively Activate T-Cells

We next evaluated the effects of E1-DHFR^2^-αCD3 against a 3D spheroid model using A431-R cells. Using live-cell imaging and Z-stacking on the Cytation C10, we show that E1-DHFR^2^-αCD3 causes spheroids to disperse over time **(Figure S4A)**. By quantifying how many fragments appear over time using the Cytation C10 cell counting software, we demonstrate that E1-DHFR^2^-αCD3 CSANs exhibit maximal disruption of spheroids over 72 hours compared to E1-DHFR^2^-αCD3 monomer and PBS control **(Figure 4D-E)**.

We then analyzed the ability of E1-DHFR^2^-αCD3 to activate T-cells by monitoring the levels of early and late T-cell activation markers CD69 and CD25, respectively. Notably, E1-DHFR^2^-αCD3 CSANs significantly increased expression of CD69 and CD25 on both CD4+ and CD8+ T-cells compared to E1-DHFR^2^-αCD3 monomer. Furthermore, T-cell activation was minimal when E1-DHFR^2^-αCD3 monomer and CSANs were incubated with T-cells and no target A431-R cells. A modest increase in CD69 on T-cells incubated with E1-DHFR^2^-αCD3 CSANs and no target cells was noted **(Figure 4F)**. This result has been seen with other bispecific T-cell engagers^17^ and can be expected due to the rapid appearance of surface CD69 after TCR/CD3 engagement^18^.

### E1-DHFR^2^-αCD3 Can Form Trifunctional CSANs and they Can be Disassembled from the Cell Surface Using Trimethoprim

It has been shown that upon ligand binding, EGFR is activated and internalized through endocytosis, trafficked to the early endosomal compartment, and either degraded or recycled to the cell surface.^16,19^ Previously, we have shown that Alexa Fluor 647 conjugated CSANs prepared from E1-DHFR^2^ monomer were able to bind EGFR+ cells and undergo rapid internalization.^11^ Additionally, we have demonstrated that bispecific CSANs can be formed by mixing different ratios of monomer X with monomer Y and that the valency of CSANs depends on the ratio of the two monomers.^9^ Consequently, we hypothesized that when the E1-DHFR^2^-αCD3 monomer is mixed with another DHFR^2^ monomer, they would potentially assemble into trifunctional CSANs, allowing their binding to EGFR+ cells and subsequent internalization to be observed.

To test our hypothesis, we prepared trifunctional E1-αCD3/green fluorescent protein (GFP) CSANs from E1-DHFR^2^-αCD3 and GFP-DHFR^2^ monomers by mixing a 1:1 ratio of the respective monomers in the presence of bis-MTX **(Figure 5A)**. The GFP-DHFR^2^ monomers consisted of DHFR^2^ and a modified GFP derived from *Aequorea Victoria* c, msGFP2, which has been reported to be less susceptible to photobloeaching.^20^ The GFP-DHFR^2^ monomer was shown to be capable of forming GFP-DHFR^2^ CSANs **(Figure S5A)**. We, therefore, assembled E1-αCD3/GFP trispecific CSANs, from a 1:1 mixture of E1-DHFR^2^-αCD3 monomer and GFP-DHFR^2^ monomer. E1-DHFR^2^-αCD3/GFP trispecific CSANs and E1/GFP bispecific CSANs were shown to bind to A431-R cells (5.8 × 10^6^ EGFR per cell) at 4 °C **(Figure 5B)**. When the temperature was raised to 37 °C, the E1-αCD3/GFP trispecific CSANs and E1/GFP bispecific CSANs were shown to undergo internalization, indicative of EGFR-mediated endocytosis as observed before by E1-DHFR^2^ CSANs.^11^

**Figure 5.**
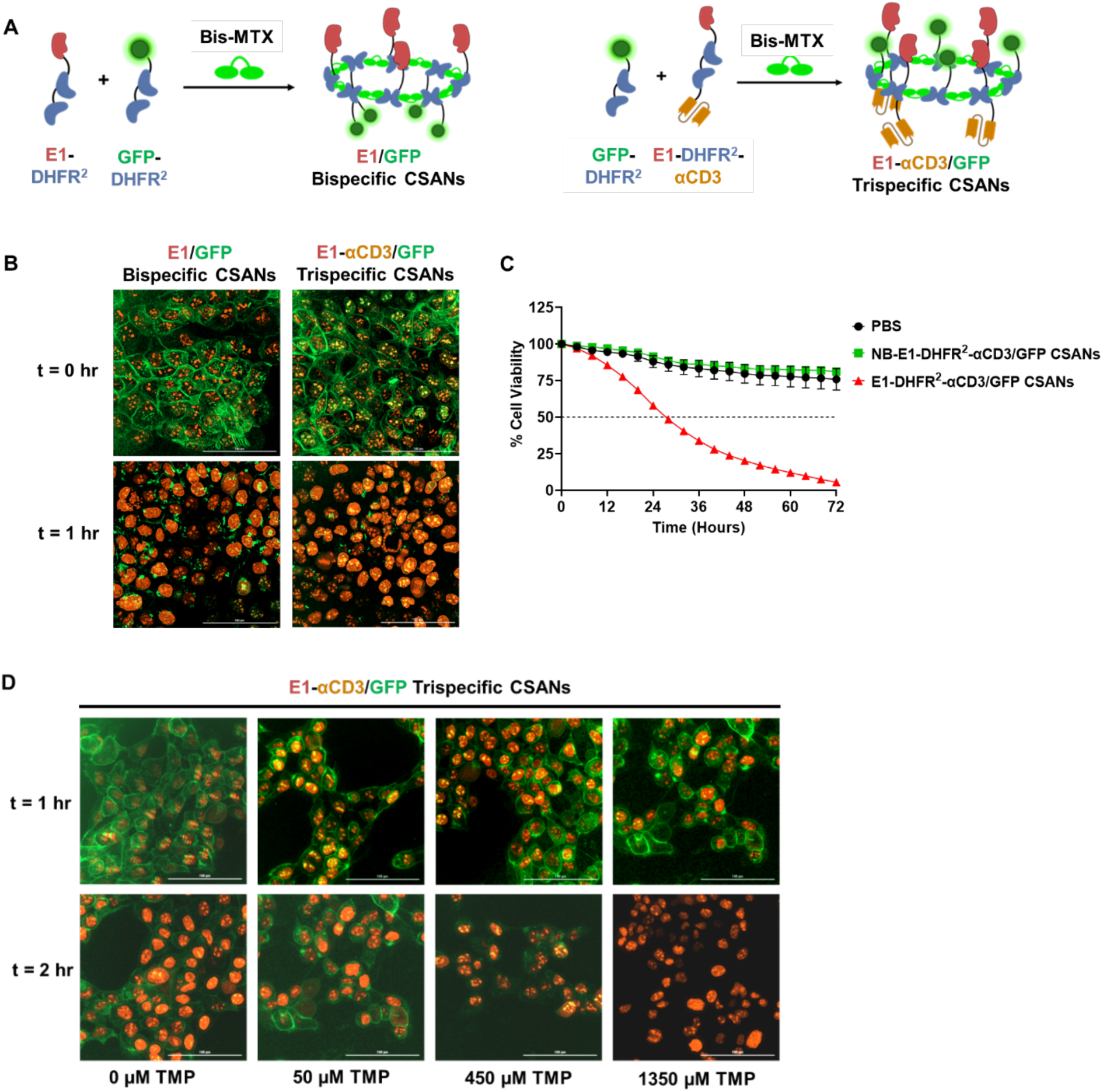
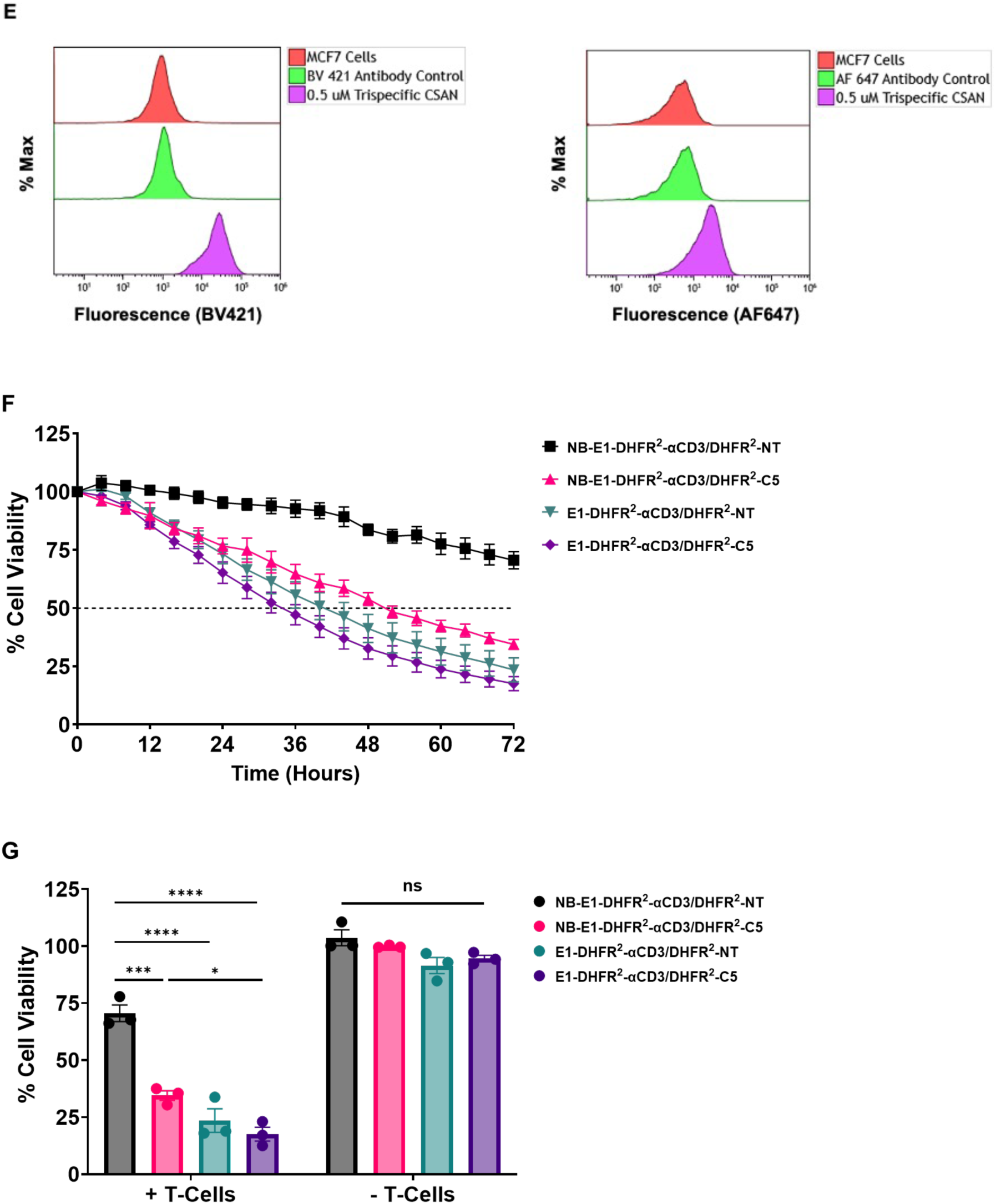
Validation, characterization, and cytotoxicity of trifunctional CSANs. **A)** CSAN formation schematic showing incorporation of a GFP-DHFR^2^ protein into either E1-DHFR^2^ or E1-DHFR^2^-αCD3 CSAN. **B)** A431-R cells were plated and allowed to adhere overnight. CSANs were added to wells and allowed to bind at 4 °C. The plate was washed to remove unbound CSANs and imaged (t=0) and then placed in 37 °C incubator. Plates were then imaged again (t=1 hour) to visualize the internalization of bound CSANs. **C)** Cytotoxicity of the E1-αCD3/GFP CSAN was evaluated. Briefly, A431-R cells were plated and allowed to adhere overnight. CSANs were prepared at a concentration of 50 nM and then added to A431-R cells with isolated donor T-cells at a ratio of 5:1 (effector-to-target). Cytotoxicity was evaluated for 72 hours using Biotek Cytation C10 at 4x magnification. The NB-E1-αCD3/GFP CSAN was included as a negative control. Data are presented as mean ± SEM of n=3 technical replicates. Error bars not shown are too small to be visualized. **D)** Disassembly of trifunctional rings with trimethoprim (TMP). CSANs were formed with equimolar ratios of E1-DHFR^2^-αCD3 and GFP-DHFR^2^. These trifunctional CSANs were then added to A431-R cells and imaged at t=1 hour. Increasing concentrations of TMP were then added at 4 °C to prevent internalization of CSANs. Cells were washed and reimaged, with the disappearance of green fluorescence indicating disassembly of rings from the surface of A431-R cells. **E)** Flow cytometry verification of trifunctional CSANs using equimolar ratios of FLAG-tagged E1-DHFR^2^-αCD3 and c-Myc-tagged DHFR^2^-C5. Briefly, MCF-7-R cells (EpCAM+ and EGFR-) were incubated with 0.5 μM E1-DHFR^2^-αCD3/DHFR^2^-C5 trispecific CSANs for 1 hour at 4 °C. Cells were washed and stained with αFLAG Brilliant Violet 421 and αc-Myc Alexa Fluor 647 antibodies. Cells were then analyzed on the Cytoflex-S cytometer. The presence of both FLAG and c-Myc tags on a single cell population indicated that both E1-DHFR^2^-αCD3 and DHFR^2^-C5 are present in the trifunctional CSANs bound to the MCF-7-R cells. **F)** Trifunctional CSANs were then evaluated for cytotoxicity against MCF-7-R cells. MCF-7-R cells were plated in a 96-well plate and allowed to adhere overnight. T-cells were isolated from human donor PBMCs and added to trifunctional CSANs for 1 hour to form PAR T-cells. The PAR T-cells were then added to MCF-7-R cells at an effector-to-target ratio of 5:1. Samples were also included with CSANs and no effector T-cells as controls. Cytotoxicity was analyzed for 72 hours by live-cell imaging with the Biotek Cytation C10. **G)** Endpoint cytotoxicity was analyzed. Data are presented as mean ± SEM of n=3 replicates with statistical significance denoted as * P<0.05, ** P<0.01, *** P<0.001, and **** P<0.0001 with ordinary one-way ANOVA followed by Tukey’s multiple comparisons test. Data obtained from T-cells isolated from a single PBMC donor but is representative of two different donors.

To verify that incorporation of GFP-DHFR^2^ did not inhibit the cytotoxic effects of E1-DHFR^2^-αCD3 *cis*-CSANs, cytotoxicity studies were carried out. The trifunctional E1-αCD3/GFP CSANs exhibited potent cytotoxicity against A431-R cells. We also prepared a point-mutant non-binding E1-DHFR^2^-αCD3 protein [herein referred to as NB-E1-DHFR^2^-αCD3] and showed that NB-E1-DHFR^2^-αCD3/GFP trispecific CSANs do not exhibit any T-cell mediated cytotoxicity towards A431-R cells **(Figure 5C)**. This further demonstrates that the CSANs cytotoxicity is specific and dependent on EGFR expression by the tumor cells.

Our lab has shown that CSANs can be disassembled from the surface of cells using the common antibiotic, trimethoprim (TMP).^13^ This serves as a versatile safety mechanism to control potential toxicities. Here we demonstrate this unique ability for the first time using trifunctional CSANs. Using the same trifunctional E1-αCD3/GFP CSANs as above, we demonstrate the dose-dependent disassembly of CSANs from the surface of A431-R cells after treatment with trimethoprim via confocal imaging **(Figure 5D)**. At the highest dose of 1350 μM trimethoprim, no green fluorescence can be seen on A431-R cells, indicating that CSANs have been successfully disassembled and removed.

### E1-DHFR^2^-αCD3 Can Form Trispecific TCE with Enhanced Cytotoxicity

Since we had shown that we can assemble trifunctional CSANs, we hypothesized that pairing E1-DHFR^2^-αCD3 with another antigen targeting DHFR^2^ monomer would form trifunctional and thus trispecific CSANs. Recently, we developed an epithelial cell adhesion molecule (EpCAM) binding fibronectin, C5, when fused to DHFR^2^ could be self-assembled into αEpCAM CSANs.^9^ EpCAM is known to be widely expressed on a wide range of epithelial solid tumors, including TNBC. Consequently, we prepared E1-αCD3/αEpCAM trispecific CSANs by treating FLAG-tagged E1-DHFR^2^-αCD3 and c-Myc-tagged DHFR^2^-C5 with bis-MTX. Flow cytometry analysis revealed that the trispecific CSANs were able to bind to cell surface expressed EGFR (5 × 10^3^ per cell) and EpCAM (1.2 × 10^6^ per cell) on MCF-7-R cells, demonstrating their trifunctionality. **(Figure 5E)**. Cytotoxicity studies demonstrated that trispecific CSANs composed of both NB-E1-DHFR^2^-αCD3 and DHFR^2^ monomer (non-targeted, NT) were unable to induce efficient T-cell mediated killing of MCF-7-R tumor cells. In contrast, CSANs composed of E1-DHFR^2^-αCD3 and DHFR^2^-NT monomers (EGFR-targeting) or NB-E1-DHFR^2^-αCD3 and DHFR^2^-C5 (EpCAM-targeting) induced similarly potent T-cell mediated MCF-7R cytotoxicity, despite a 1000-fold difference in antigen expression between EGFR and EpCAM **(Figure 5F-G)**. Moreover, trispecific CSANs composed of E1-DHFR^2^-αCD3 and DHFR^2^-C5 (EGFR and EpCAM-targeting) were shown to have enhanced potency over CSANs composed of NB-E1-DHFR^2^-αCD3 and DHFR^2^-C5, indicating that the targeting of EGFR augmented EpCAM targeting by the E1-αCD3/C5 trispecific CSANs.

## Discussion

T-cell engagers have emerged as a viable strategy for targeted cancer immunotherapy.^4^ However, success against solid tumors has been limited, as well as durable responses. To enhance efficacy, combination approaches have been investigated by pairing TCEs against multiple targets, such as two different tumor antigens or immune checkpoints, like PD-L1.^21^ In this study, we designed and characterized a self-assembling protein nanoring-based bispecific T-cell engager referred to as chemically self-assembled nanorings (CSAN) that enables multivalent, trifunctional targeting. By engineering the DHFR^2^ protein with both an anti-EGFR fibronectin (E1) and an anti-CD3 scFv, we successfully created a bifunctional E1-DHFR^2^-αCD3 monomer capable of binding both T-cells and EGFR-expressing tumor cells. Upon self-assembly with bis-MTX, the resulting *cis*-CSANs exhibited a smaller hydrodynamic diameter compared to traditional *trans*-CSANs, suggesting that the *cis* configuration influences CSAN assembly and possibly enhances ring compactness with reduced valency. The ability to prepare bispecific multivalent CSANs from the self-assembly of a single protein is an advantage over previous approaches, in which two protein monomers with differing specificities have been used.^10–12^ Additionally, the *cis*-CSANs approach enables a direct comparison between a mono-valent bispecific TCE and a multi-valent bispecific TCE.

The E1-DHFR^2^-αCD3 monomer and its corresponding *cis*-CSANs were shown to facilitate bifunctional binding and, similar to previous EGFR-targeting CSANs, undergo internalization by EGFR+ cells. Additionally, potent T-cell-mediated cytotoxicity was observed against multiple EGFR+ tumor cell lines. T-cell mediated destruction of 3D spheroids was also demonstrated using novel live-cell imaging and 3D spheroid techniques. As expected for TCE, the cytotoxicity of the CSANs was shown to be both concentration- and effector-to-target (E:T) ratio-dependent. In addition, the kinetics and level of cytotoxic response were found to be strictly dependent on both the presence of EGFR expression and T-cells, indicating a high degree of target specificity. Moreover, comparable cytotoxic efficacy was observed across T-cells from multiple donors, highlighting the potential for robust and consistent therapeutic effects across different patient populations.

Building on the versatility of the E1-DHFR^2^-αCD3 monomer, we further demonstrated that it could be used to prepare trifunctional CSANs by co-assembling with additional DHFR^2^ fusion monomers. We generated E1-αCD3/GFP and E1-αCD3/C5 trispecific CSANs that maintained functional binding, induced internalization, and cytotoxicity. Strikingly, trispecific CSANs targeting both EGFR and EpCAM demonstrated enhanced cytotoxicity compared to bispecific CSANs targeting just EpCAM, suggesting that multi-specific CSANs could mitigate issues like tumor antigen heterogeneity and antigen escape—major hurdles in solid tumor immunotherapy.^22,23^ Taken together, these results highlight that bispecific monomers, such as E1-DHFR^2^-αCD3, can not only form potent bispecific *cis*-CSAN-TCEs but also serve as a modular platform to create customizable trifunctional and trispecific TCEs. Thus, CSANs could be easily adapted to target different combinations of tumor-associated antigens, enhancing the specificity and breadth of TCEs, while potentially incorporating fluorophores and immune activators.

## Summary

We have demonstrated that a bifunctional E1-DHFR2-αCD3 fusion protein can be used to form potent bispecific multivalent *cis*-CSANs capable of inducing targeted T-cell mediated killing of EGFR+ tumor cells. These bispecific *cis*-CSANs maintain high specificity, robust cytotoxicity, and donor-independent activity in both 2D and 3D spheroid models. Furthermore, we showed that CSANs, self-assembled from E1-DHFR^2^-αCD3 and other labeled or targeting DHFR^2^ monomers, retain functional activity and exhibit enhanced cytotoxicity. This work establishes CSANs as a versatile and modular platform for evaluating the therapeutic potential of multi-specific TCE to address tumor cellular antigen heterogeneity and immune evasion.

## Acknowledgements

We gratefully acknowledge support from NCI R01CA247681 (CRW), a traineeship to BM from NIGMS -T32 GM008700 and a NSF Graduate Fellowship to BM.

## Declaration of generative AI and AI-assisted technologies in the writing process

During the preparation of this work the corresponding author used Grammerly in order to assist in correcting typing and textual mistakes. After using this tool, the authors reviewed and edited the content as needed and take full responsibility for the content of the publication.

## Methods

### Cell Lines, Culture Conditions, and T-Cell Isolation

A431, MDA-MB-231, and MDA-MB-436 were obtained from the American Type Culture Collection (ATCC, Rockville, MD). To produce fluorescent stable cell lines, cells were transduced with either Incucyte® NucLight green or red lentivirus and selected with puromycin. Cells were cultured in Dulbecco’s Modified Eagle Medium (DMEM) supplemented with 10% fetal bovine serum (FBS) and 1% Thermo Fischer (Waltham, MA) GlutaMAX at 37 °C in 5% CO_2_. Human peripheral blood mononuclear cells (PBMCs) from healthy donors were isolated from buffy coats (obtained from Memorial Blood Centers, St. Paul, MN) by density gradient centrifugation using Greiner Bio-One Leucosep centrifuge tubes. T-Cells were subsequently isolated from PBMCs using Akadeum Life Sciences (Ann Arbor, MI) Human T-Cell Isolation Kit.

### EGFR Antigen Expression Quantification

EGFR expression was quantified using Bangs Laboratory QSC Anti-Mouse IgG Fluorescence Calibration Beads (Fischer, IN). Cells and beads were incubated with Brilliant Violet 421 anti-human EGFR antibody (BioLegend, Cat: 352911) for 1 hour, washed, and analyzed using Cytoflex-S flow cytometer (Beckman Coulter) and Kaluza analysis software (Beckman Coulter).

### DHFR^2^ Protein Expression and Purification

For EGFR targeting, an EGFR-binding fibronectin (clone E1)^11^ was fused to the N-terminus of the DHFR^2^ construct separated by a glycine-serine linker. For T-cell targeting, a previously designed aCD3-scFv derived from monoclonal antibody, UCHT1, was fused to the C-terminus of the DHFR^2^ construct. A single glycine linker was used between the two DHFRs. A 15 amino acid glycine-serine linker was used between the C-terminal DHFR and the αCD3-scFv. The complete E1-DHFR^2^-αCD3 plasmid was transformed into T7 Express cells from New England Biolabs (Ipswich, MA). Colonies picked were cultured in 1 liter of 2XYT media, shaken at 180 rpm at 37 °C until OD_600_ value reached 0.6, and induced with 1mM final concentration of isopropyl b-D-1-thiogalactopyrranoside (IPTG; Sigma-Aldrich, St. Louis, MO). Cultures were incubated for 18 hours at room temperature and pelleted for inclusion body purification as described before.^6^

For GFP-DHFR^2^, the msGFP2 sequence was fused to the N-terminus of the DHFR^2^ construct separated by a glycine-glycine-glycine linker. An N-terminal His tag was included for purification and detection. The GFP-DHFR^2^ plasmid was transformed into T7 Express cells. Colonies picked were cultured in 1 liter of 2XYT media, shaken at 180 rpm at 37° C until OD_600_ value reached 0.6, and induced with 1mM final concentration of isopropyl b-D-1-thiogalactopyrranoside (IPTG; Sigma-Aldrich, St. Louis, MO). Cultures were incubated for 4 hours at 37° C, pelleted, and lysed. The soluble fraction was collected and purified using Cytiva Talon® Superflow™ histidine-tagged protein purification resin (Marlborough, MA).

### CSAN Oligomerization and Characterization

CSANs were formed by mixing E1-DHFR^2^-αCD3 monomer with 5 equivalents of the chemical dimerizer, bis-methotrexate. The solution was incubated for 1 hour at 37° C prior to characterization by analytical SEC. Samples were also analyzed by dynamic light scattering (DLS) using Anton Paar Litesizer 500 (Ashland, VA). Hydrodynamic diameter of E1-DHFR^2^-αCD3 monomer and CSANs are shown as mean ± SD of three replicates.

### Binding Assays

To characterize EGFR binding, A431 cells were trypsinized and 1.0 X 10^5^ cells were incubated with E1-DHFR^2^-αCD3 monomer and CSANs for 1 h at 4° C. Cells were washed with FACS buffer and incubated with anti-FLAG phycoerythrin (PE) conjugated antibody (BioLegend clone L5) for 45 minutes in the dark at 4 °C. Cells were washed again with FACS buffer and analyzed using Cytoflex-S flow cytometer and Kaluza software. Jurkat T-cells were used to characterize CD3 binding, and samples were prepared identically to above. Median fluorescence intensity (MFI) was monitored and compared to unstained and control samples.

### Internalization and Disassembly Assays

Trispecific CSANs were prepared by mixing equimolar amounts of E1-DHFR^2^-αCD3 with GFP-DHFR^2^ in the presence of bis-methotrexate and incubating for 1 hour at 37 °C. CSANs were added to A431-R cells cultured overnight in 96-well Agilent (Santa Clara, CA) flat-bottom plates and incubated for 1 hour at 4 °C. Cells were then washed with PBS and imaged at 60x magnification using the Agilent BioTek Cytation C10 Confocal Imager. For internalization studies, plates were then incubated for 1 hour at 37 °C prior to washing and imaging. For disassembly assays, cells were incubated with varying concentrations of trimethoprim at 4 °C for 1 hour prior to washing and imaging.

### 2D Cytotoxicity Assays

Fluorescent cancer cells (constitutively expressing either GFP or mKate2) were plated at 5.0 X 10^3^ cells/well in Agilent 96-well flat-bottom plates and rested overnight. T-cells isolated and expanded from healthy donor PBMCs were thawed and rested overnight in Roswell Park Memorial Institute (RPMI) medium supplemented with 10% FBS and 1% Thermo Fischer GlutaMAX. The next day, T-cells were functionalized with CSANs for 1 hour and added to wells containing target cells (n=3 wells per treatment). Plates were imaged every 4 hours using the Agilent BioTek Cytation C10 Confocal Imager at 4x magnification for 72 hours. Fluorescent object area was quantified using BioTek Gen5 software (Agilent) and data is shown as mean ± SEM. Data is shown as a ratio of treated/untreated cells fluorescent object area. Data was analyzed using GraphPad Software (Boston, MA).

### 3D#Cytotoxicity Assays

5.0 X 10^3^ A431-R cells were seeded in Agilent 96-well ULA round-bottom plates. Plates were centrifuged at 180 rpm for 10 minutes and spheroids were allowed to form overnight. T-cells were revived as described above. The next day, T-cells were functionalized with 50 nM CSANs for 1 hour and added to wells containing spheroids (n=3 wells per treatment). Plates were imaged every 4 hours using the Agilent BioTek Cytation C10 Confocal Imager at 20x magnification for 72 hours. BioTek Gen5 software was used to create Z-projections of all slices of spheroids and spheroid area of these Z-projections was quantified. To quantify spheroid destruction a percent disruption was calculated using the cell masking function to track the number of pieces in the spheroid as they were broken apart by CSAN treatment. A higher percent disruption indicates a higher amount of spheroid destruction.

### T-Cell Activation Assay

1.0 X 10^4^ A431 cells were seeded in Agilent 96-well flat-bottom plates and rested overnight. PBMCs were revived and rested overnight in RPMI supplemented with 10% FBS and 1% Thermo Fischer GlutaMAX. The next day, T-cells were isolated using the Akadeum Life Sciences Human T-Cell Isolation Kit. 1.0 X 10^5^ T-cells were added to wells with 25 nM E1-DHFR^2^-αCD3 monomer or CSANs either in the presence or absence of target cells and incubated at 37 °C for 24 hours. T-cells were then collected and stained with PE anti-human CD4 (BioLegend, Cat: 980804), Brilliant Violet™ 750 anti-human CD8 (BioLegend, Cat: 344755), FITC anti-human CD25 (BioLegend, Cat: 302603), and APC anti-human CD69 (BioLegend, Cat: 985206). Samples were run on the Cytoflex-S flow cytometer and analyzed with Kaluza software.

---

**Supplemental Figure 1.**
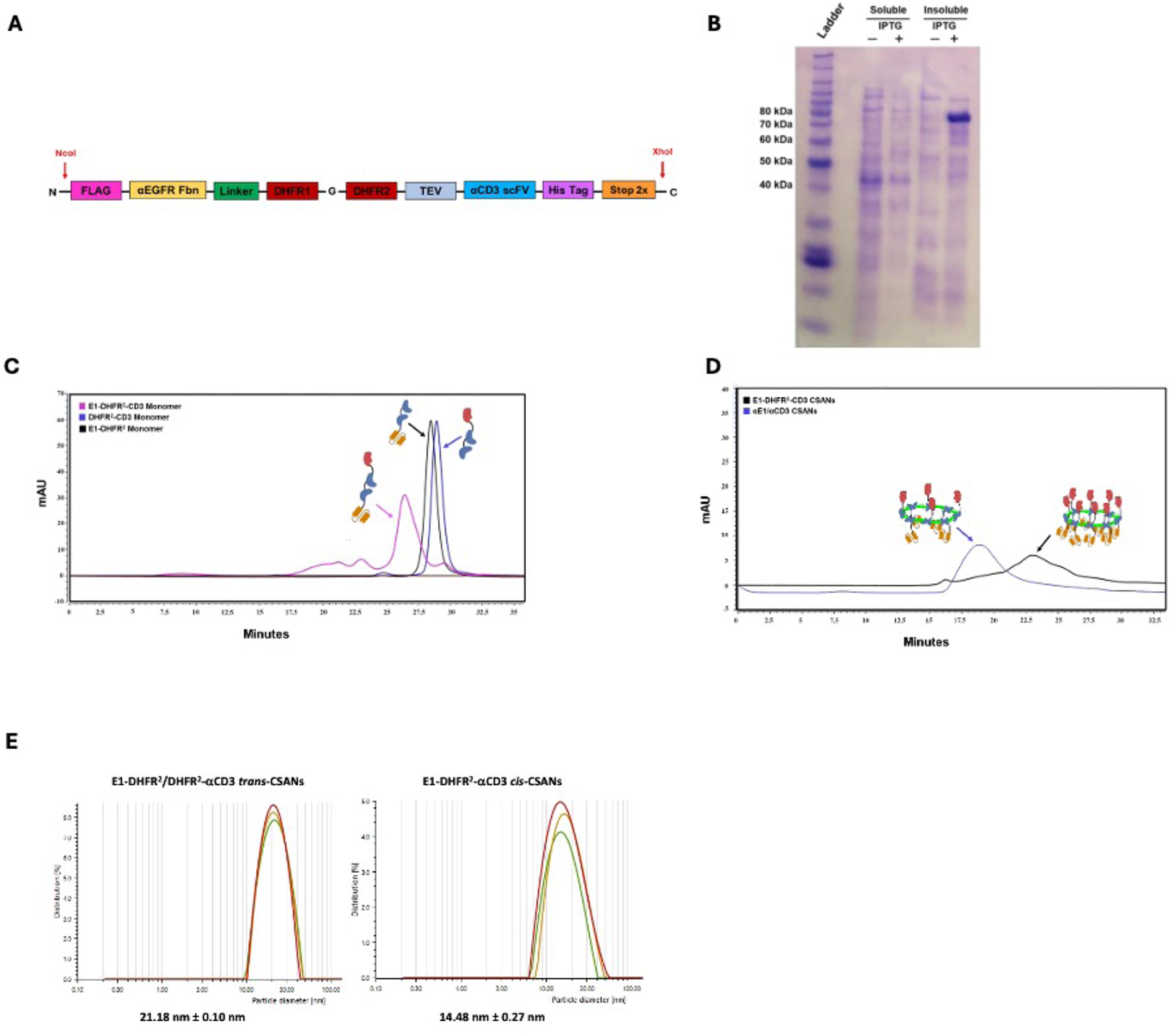
Design, characterization, and comparison of E1-DHFR^2^-αCD3 construct to *trans* E1-DHFR^2^/DHFR^2^-αCD3 CSANs. **A)** Sequence for E1-DHFR^2^-αCD3 was designed as a gblock and cloned into pET28 vector using NcoI and XhoI restriction enzymes. **B)** SDS-PAGE analysis confirms successful induction of E1-DHFR^2^-αCD3 which has a MW of 78.5 kDa. **C)** Confirmation of monomeric E1-DHFR^2^-αCD3 protein using SEC analysis showing larger size (earlier elution) compared to monomeric components of *trans* CSANs (E1-DHFR^2^ and DHFR^2^-αCD3). **D)** SEC comparison of ring formation for *cis* vs *trans* CSANs which show a paradoxical smaller ring size for E1-DHFR^2^-αCD3 *cis* CSANs. **E)** Confirmation of smaller E1-DHFR^2^-αCD3 *cis* CSAN ring size using dynamic light scattering when compared to *trans* CSANs. Data represent average of 3 separate runs ± SEM.

**Supplemental Figure 2.**
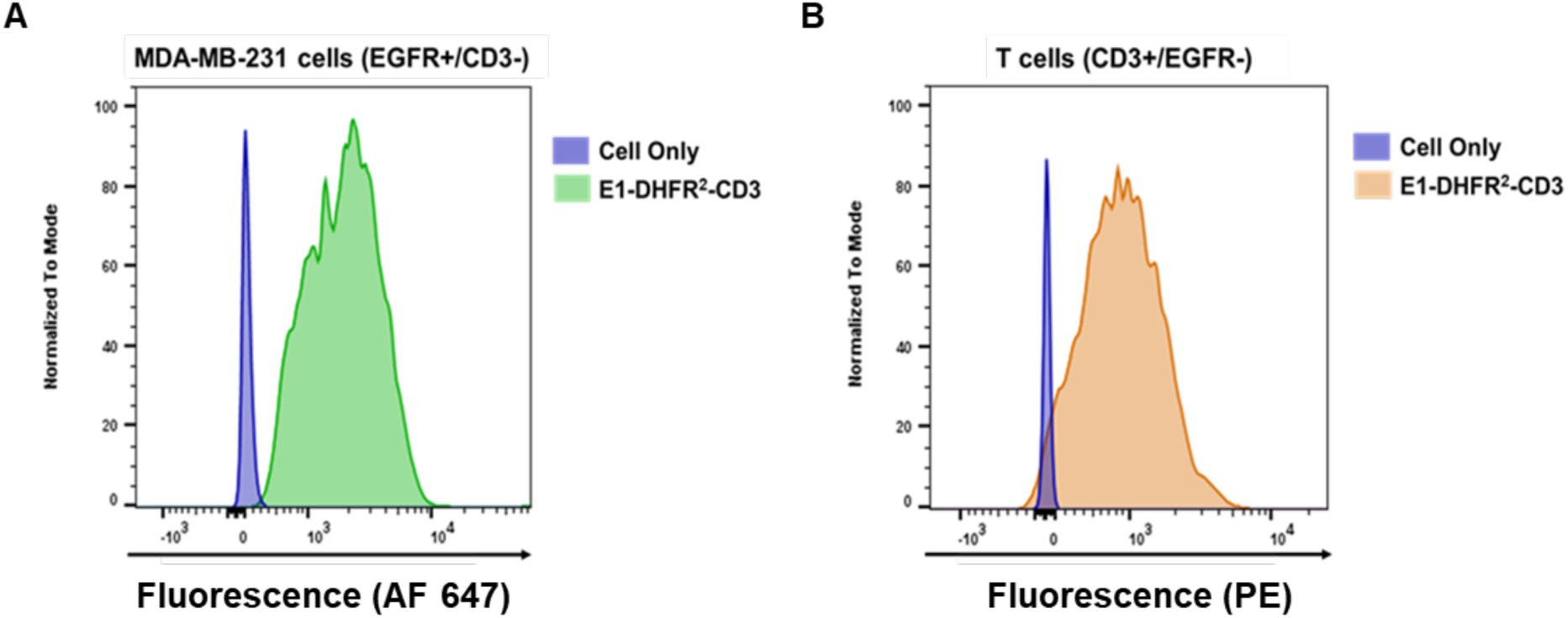
Flow cytometry validation of E1-DHFR^2^-αCD3 monomer binding. **A)** E1-DHFR^2^-αCD3 functionality confirmed via binding to EGFR+/CD3- MDA-MB-231 cells and **B)** CD3+/EGFR- human T-cells by flow cytometry.

**Supplemental Figure 3.**
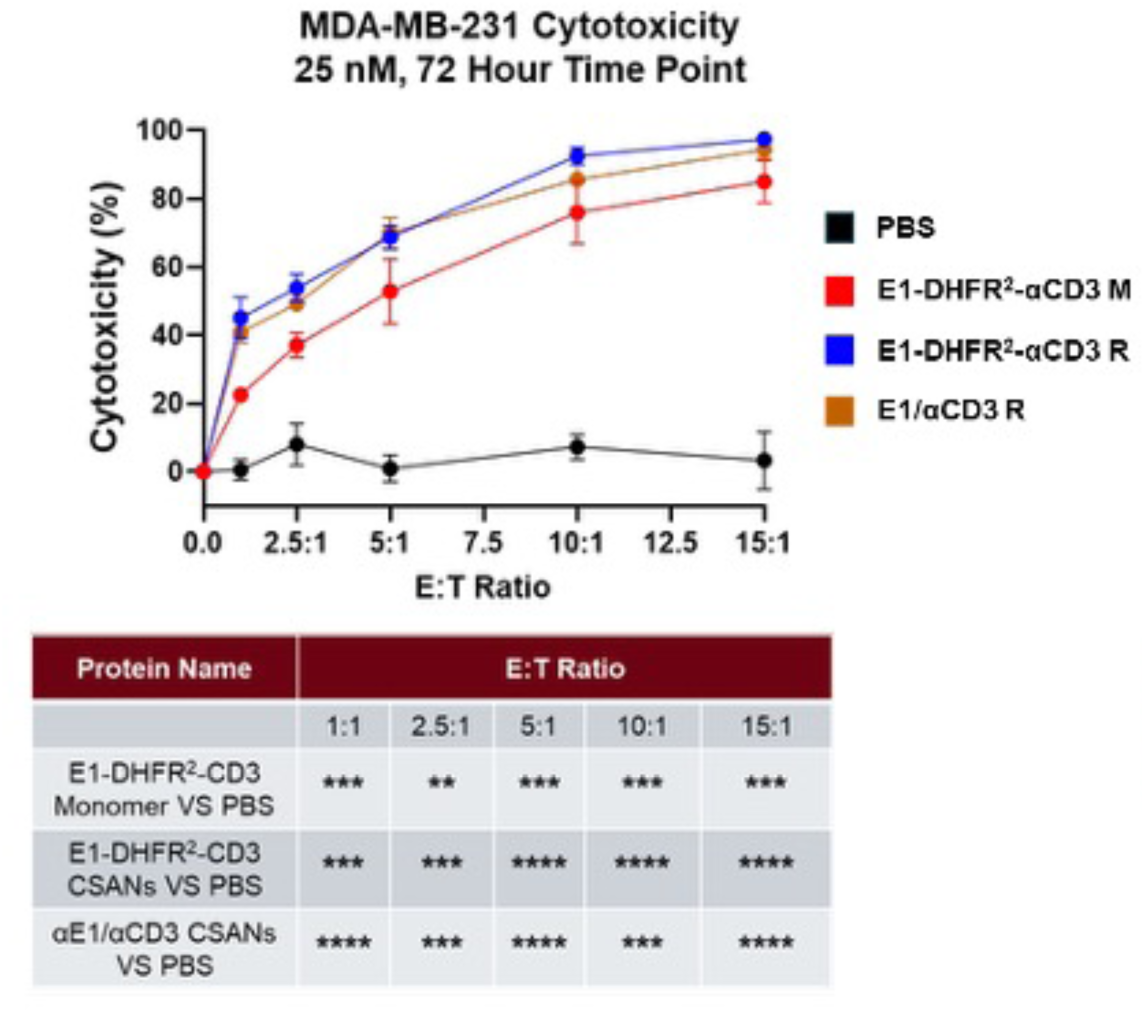
Multiple E:T ratio validation of E1-DHFR^2^-αCD3 monomer and *cis-* CSAN cytotoxicity against MDA-MB-231-G cells. T cells isolated from donor-derived PBMCs were modified with E1-DHFR^2^-αCD3 monomer, E1-DHFR^2^-αCD3 *cis-*CSANs, and E1-DHFR^2^/DHFR^2^-αCD3 *trans* CSANs at 25 nM to generate PAR T cells, which were then introduced to the cell monolayer at varied E:T ratios. Cell viability was measured over 72 hours. Data are presented as mean ± SEM with statistical significance denoted as * P<0.05, ** P<0.01, and *** P<0.001 with respect to controls by two-way ANOVA followed by Dunnett’s test for multiple comparisons. Data were obtained from a single donor, where n=3 replicates.

**Supplemental Figure 4-3D spheroid timelapse imaging**

See PPT file-*“Supplemental figure 4A.pptx”*

**Supplemental Figure 5.**
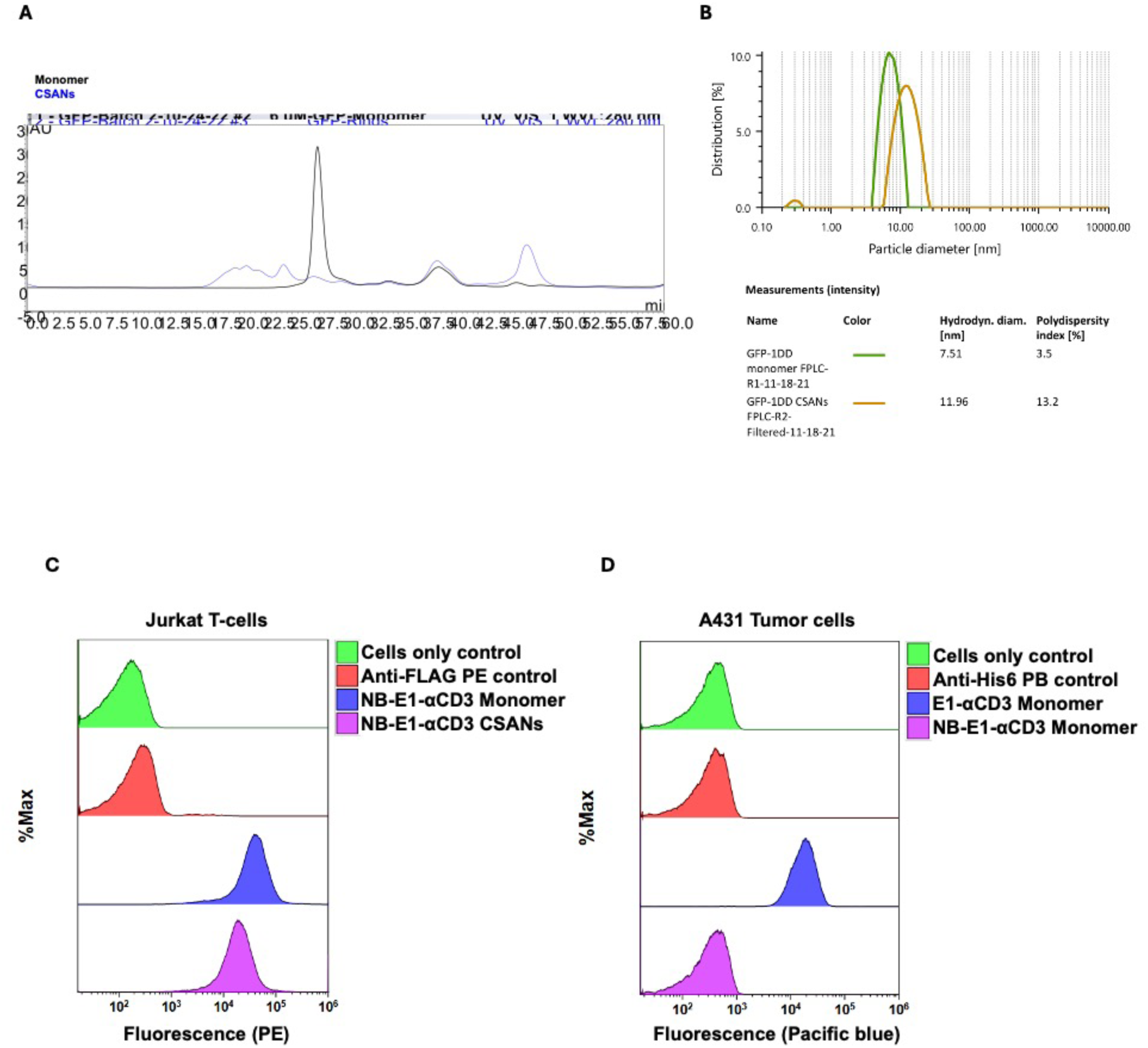
GFP-DHFR^2^ and NB-E1-DHFR^2^-αCD3 characterization. **A)** SEC trace of GFP-DHFR^2^ monomer and CSAN demonstrating purity and ability to oligomerize after incubation with bis-MTX. **B)** Dynamic light scattering (DLS) of GFP-DHFR^2^ monomer and CSANs demonstrating purity and increased hydrodynamic radius of GFP-DHFR^2^ CSANs after 1 hour incubation with bis-MTX. **C)** SEC trace confirming purity of NB-E1-DHFR^2^-αCD3 after refolding. **C)** Flow cytometry validation of NB-E1-DHFR^2^-αCD3 binding characteristics. NB-E1-DHFR^2^-αCD3. Both NB-E1-DHFR^2^-αCD3 ring and monomer bound to Jurkat T-cells (CD3+). **D)** NB-E1-DHFR^2^-αCD3 did not bind to EGFR+ A431 cells compared to αEGFR (E1)-DHFR^2^-αCD3.

**Supplemental Figure 6.**
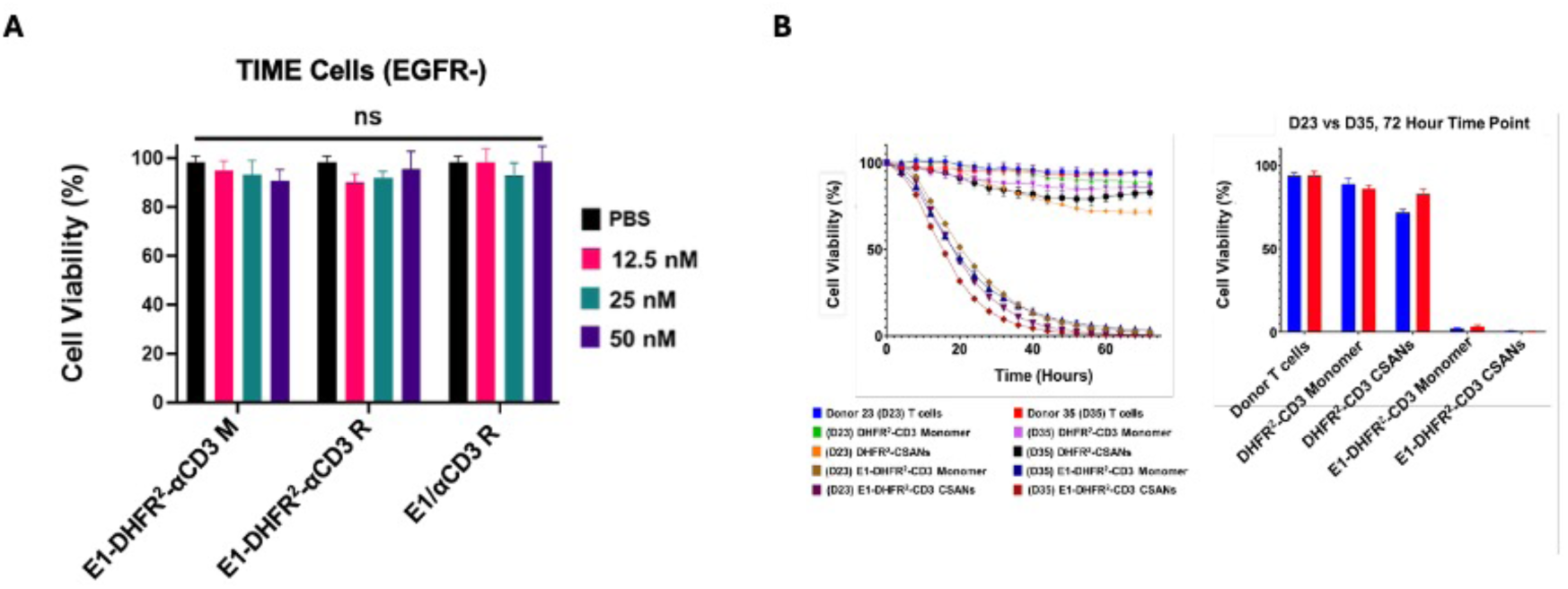
Evaluation of cytotoxicity on EGFR-cell line and multidonor cytotoxicity validation. **A)** EGFR negative TIME cells were seeded into a 96 well plate and incubated with all three proteins alongside isolated T-cells from donor derived PBMCs. Cytotoxicity was evaluated after 72 hours. Data are presented as mean ± SEM with statistical significance denoted as (* P<0.05, **P<0.01, ***P<0.001, and ****P<0.0001 with respect to the PBS control by two-way ANOVA followed by Dunnett’s test for multiple comparisons. Data were obtained from a single donor, which n = 3-9 biological replicates, each with technical triplicates.) **B)** To ensure reproducibility, T-cells were isolated from two different human PBMC donors. A431 cells were grown as a monolayer in a 96-well plate. T-cells from two donors were added to appropriate cells, followed by the addition of E1-DHFR^2^-αCD3 monomer and CSANs (targeted), and DHFR^2^-αCD3 monomer and CSANs (non-targeted). Cytotoxicity was assessed for both donors after 72 hours. Data from both experiments are presented as mean ± SEM with n=3 replicates.

**Supplemental Table 1.** Cell line receptor expression of EGFR and EpCAM. Cell line receptor expression of EGFR and EpCAM were characterized by incubating cell with Brilliant Violet 421 anti-human EGFR antibody (BioLegend, Cat: 352911) or PE anti-human EpCAM antibody (Biolegend, Cat: 324305). Receptor expression was quantified using Bangs Laboratory QSC Anti-Mouse IgG Fluorescence Calibration Beads (Fischer, IN). N.D.= Not detected.

| Cell Line | EGFR<br>Receptor per cell<br>(% *Expression*) | EpCAM<br>Receptor per cell<br>(% *Expression*) |
| --- | --- | --- |
| A431 | 5.8x10<sup>6</sup> (100%) | 2.6x10<sup>5</sup> (99%) |
| MCF-7 | 5x10<sup>3</sup> (9.5%) | 1.2x10<sup>6</sup> (100%) |
| MDA-MB-231 | 6.8x10<sup>4</sup> (100%) | 8.8x10<sup>3</sup> (99%) |
| MDA-MB-436 | 3.9x10<sup>4</sup> (98%) | N.D. |

